# ModelTeller: model selection for optimal phylogenetic reconstruction using machine learning

**DOI:** 10.1101/2020.01.09.899906

**Authors:** Shiran Abadi, Oren Avram, Saharon Rosset, Tal Pupko, Itay Mayrose

**Affiliations:** School of Plant Sciences and Food security, Tel Aviv University, Ramat Aviv, Tel-Aviv 69978, Israel; School of Molecular Cell Biology & Biotechnology, Tel Aviv University, Ramat Aviv, Tel-Aviv 69978, Israel; Department of Statistics and Operations Research, School of Mathematical Sciences, Tel Aviv University, Ramat Aviv, Tel-Aviv 69978, Israel

**Keywords:** model selection, phylogenetic reconstruction, simulations, nucleotide substitution models, machine learning, Random Forest for regression

## Abstract

Statistical criteria have long been the standard for selecting the best model for phylogenetic reconstruction and downstream statistical inference. While model selection is regarded as a fundamental step in phylogenetics, existing methods for this task consume computational resources for long processing time, they are not always feasible, and sometimes depend on preliminary assumptions which do not hold for sequence data. Moreover, while these methods are dedicated to revealing the processes that underlie the sequence data, in most cases they do not produce the most accurate trees. Notably, phylogeny reconstruction consists of two related tasks, topology reconstruction and branch-length estimation. It was previously shown that in many cases the most complex model, GTR+I+G, leads to topologies that are as accurate as using existing model selection criteria, but overestimates branch lengths. Here, we present ModelTeller, a computational methodology for phylogenetic model selection, devised within the machine-learning framework, optimized to predict the most accurate model for branch-length estimation accuracy. ModelTeller relies on a readily implemented machine-learning model and thus the prediction according to features extracted from the sequence data results in a substantial decrease in running time compared to existing strategies. We show that on datasets simulated under simple homogenous substitution models ModelTeller leads to branch-length estimation that is as accurate as the statistical model selection criteria. We then demonstrate that ModelTeller outperforms these criteria when more intricate patterns – that aim at mimicking realistic processes – are considered.

## Introduction

The abundance of substitution models (Jukes and Cantor 1969; Kimura 1980; Felsenstein 1981; Cowan 1984; Hasegawa et al. 1985; Tamura 1992; Tamura and Nei 1993; Schöniger and Von Haeseler 1994; Zharkikh 1994; Huelsenbeck and Crandall 1997) and the need to choose one (or few) has established model selection as a pre-requisite for phylogeny reconstruction (Goldman 1993a; Huelsenbeck and Rannala 1997; Sullivan and Swofford 1997; Posada and Crandall 2001; Sullivan and Swofford 2001; Pupko et al. 2002). This is evident by the wide use of model selection as an inherent component of phylogenetic analysis. For example, the most widely used method, MODELTEST (Posada and Crandall 1998), was included in the 100 all-time top-cited papers by Web of Science (Van Noorden et al. 2014). However, while criteria for phylogenetic model selection have been adapted from the general statistical literature, they rely on assumptions that do not hold for phylogenetic data analysis (Posada and Buckley 2004). For example, the Likelihood Ratio Test (LRT), a chi-square test between a pair of nested models, has been derived to establish the hierarchical Likelihood Ratio Tests (hLRT) criterion which performs a sequence of LRTs between pairs of nested substitution models, until a model that cannot be rejected is reached. However, LRTs assume that at least one of the compared models is adequate and might be incorrect when the models are misspecified (Foutz and Srivastava 1977; Kent 1982; Golden 1995). Obviously, no substitution model can fully capture the genuine complexity of the evolutionary process, such that even the most adequate one merely provides an approximation of reality (Box 1976), therefore posing model misspecifications that may bias the results of LRTs in phylogenetics (Zhang 1999). Furthermore, it has been shown that the choices determined by the hLRT are subject to the order of pairwise tests and the level of significance (Yang et al. 1995; Ripplinger and Sullivan 2008). Other information criteria compute the maximum-likelihood scores for all the candidate models simultaneously. As the maximum-likelihood score generally increases with the inclusion of more parameters, the Akaike and Bayesian Information Criteria, AIC (Akaike 1973; Akaike 1974) and BIC (Schwarz 1978), assign different penalties according to the number of parameters included in the model while also considering the data size. AIC assumes that the sample size is large enough to maintain the asymptotic property of the likelihood function and thus penalizes the likelihood score only for the number of model parameters. This assumption, however, rarely holds for phylogenetic data. In contrast, the corrected AIC, AICc (Sugiura 1978; Hurvich and Tsai 1989) and BIC criteria penalize the likelihood score by the number of parameters as well as the data size. Unfortunately, data size is not well defined for phylogenetic data because—owing to shared evolutionary history as well as functional and structural constraints—aligned sequences are surely dependent, as are the sites along the alignment. Although previous studies advocated to use the alignment length as the sample size (Posada and Crandall 2001; Minin et al. 2003; Posada 2008; Ripplinger and Sullivan 2008; Darriba et al. 2012), much debate exists regarding the effective data size in phylogenetics (Churchill et al. 1992; Goldman 1998; Morozov et al. 2000; Posada and Buckley 2004).

It was previously reported that model selection adds little to the performance of specific inferential tasks. For example, we and others have demonstrated that employing the most parameter-rich model, GTR+I+G, leads to topology reconstructions that are as accurate as those produced by established model selection strategies (Arbiza et al. 2011; Abadi et al. 2019; Spielman 2019). On the other hand, for the tasks of branch-length and divergence time estimations it was shown that the accuracy is conditioned on the selected model and so model selection should still be advocated (Buckley et al. 2001; Posada 2001; Minin et al. 2003; Abdo et al. 2005; Spielman and Kosakovsky Pond 2018; Dornburg et al. 2019). For example, Abadi et al. (Abadi et al. 2019) demonstrated that the consistent use of GTR+I+G generally results in inferior branch-length estimates compared to those obtained by model selection criteria, with the BIC and DT criteria obtaining the most accurate estimates. Notwithstanding, current practices focus at revealing the best-fitted model, i.e., the model that best explains the underlying process, rather than the one that produces the most accurate phylogeny. Thus, a method that is optimized to select the best model for phylogeny reconstruction should improve results (Sanderson and Kim 2000; Kelchner and Thomas 2007). Here, we employ the emerging machine-learning computational framework to tackle this task.

A supervised machine-learning technique consists of training an algorithm over descriptive datasets to learn the effect of the data characteristics on the outcome. In practice, a data sample is represented by a set of explanatory features. According to these features, an algorithm is trained to predict the most appropriate label in the case of classification, or a continuous target value in the case of regression. For example, decision trees are constructed such that every node splits the training data according to a threshold value of a certain feature (Lori and Oded 2008). Essentially, the features along paths of the tree repeatedly separate the data to smaller subsets such that the samples within a subset at a tip own similar features values beneficial to characterizing it with a certain label. A successful reconstruction of a decision tree is represented by a categorization of instances of the same label with high accuracy, and indicates the usage of powerful features. Numerous algorithms for classification or regression exist, e.g., K-means, Naïve Bayes, support vector machine (SVM), and K-Nearest-Neighbors (Kotsiantis 2007). High performance of a method for a certain dataset and learning task does not guarantee that it would perform well for another, thus the suitability of alternative algorithms should preferably be examined (Caruana and Niculescu-Mizil 2006). In any case, supplying the algorithm with a training set that extends over the full range of realistic instances is crucial for ultimate modelling of possible occurrences.

## New Approaches

We propose the use of machine learning as a novel approach for phylogenetic model selection. We present ModelTeller, which is based on the Random Forest learning algorithm, for the prediction of the optimal model for branch-length estimation. ModelTeller is trained over datasets that were simulated using parameter estimates that were drawn from an extensive cohort of empirical phylogenetic datasets, thus representing realistic data characteristics. When examined over relatively simple datasets (i.e., simulated using a single time-homogeneous model) we demonstrate that ModelTeller is as accurate as AIC and BIC in branch-lengths estimation, but with significant improvement in running time. However, such simulations are simplification of reality and do not actually represent true evolutionary processes. On a more complex simulation set we demonstrate that ModelTeller is more accurate for branch-lengths estimation than AIC and BIC. We also demonstrate that even the best models for branch-lengths estimation do not always produce topologies that are as accurate as using the most complex model, GTR+I+G. Therefore, we suggest a two-step reconstruction for optimal phylogeny reconstruction, in which topologies are first inferred by optimizing a maximum-likelihood phylogeny using the GTR+I+G model and then the best model for branch-length estimation is predicted given the GTR+I+G topology.

## Results

### Existing methodologies for phylogeny reconstruction

We first analyzed the accuracy of phylogeny reconstruction by existing methods for model selection. A total of 10,832 empirical alignments were assembled from several sequence databases. The empirical data were used to simulate realistic alignments, such that for each dataset a model was selected randomly (out of 24 commonly used nucleotide substitution models), the parameters of that model were estimated for the dataset, and in turn were used to simulate an alignment (referred to as the ‘single-model simulations’). Then, AIC and BIC were executed to select the best model for each simulated dataset. While we expected that successfully selecting the true model out of 24 nested models would be a difficult task, BIC and AIC recovered the models that generated the data for 75% and 69% datasets, respectively. However, in most phylogenetic analyses the generating model can be considered as a nuisance parameter while the accuracy of the resulting phylogeny is of utmost importance. To examine the accuracy of phylogenetic reconstruction, we reconstructed the maximum-likelihood tree according to each of the 24 models, and measured which one is more similar to the true tree when branch lengths are examined using the Branch-Score (BS) distance (Kuhner and Felsenstein 1994). In 84% and 87% of the datasets, alternative models led to branch-lengths estimates that are better than the ones inferred by the models selected by AIC and BIC, respectively. In many cases the selected models were not even the second best, namely, AIC and BIC selected the models that were ranked 8.44 and 8.24 on average, respectively (in a ranking of 1 to 24 from lowest to highest BS distances). These results indicate that although AIC and BIC select the generating models with high frequencies, these selections often yield sub-optimal branch-length estimates. Indeed, in 92% of the datasets the generating models resulted in less accurate estimates than those of alternative models, with an average rank of 8.27, again indicating that the generating model is not the optimal model for branch-length estimation.

Next, we examined whether the most complex model, GTR+I+G, can be used as a fixed model instead of applying a model selection procedure. To this end, we compared the branch-lengths estimation and topologies accuracy measurements between three reconstruction strategies: AIC, BIC, and a consistent reconstruction with the GTR+I+G model. When the accuracy of topology inference was examined for the single-model simulations, the three strategies performed similarly. GTR+I+G was only marginally inferior to AIC and BIC, but these results were statistically non-significant (Supplementary Table S1). Since these datasets were generated using a single model, thereby assuming the same evolutionary process along all parts of the phylogeny and across all positions to generate the evolutionary processes along the sequences, they do not realistically represent the complex patterns observed within empirical data. To this end, we generated additional simulated data that mimic empirical data more realistically by using more complex simulation patterns for the same empirical data. To generate the complex-model simulation set, multiple models were estimated across the alignment along with rate-variation across sites that resemble empirical data more closely than using the gamma distribution. When the topological accuracy was examined over the complex-model simulations, GTR+I+G performed significantly better than both AIC and BIC (*P = 10*^*-7*^ *and 10*^*-15*^, respectively, Wilcoxon signed rank tests). Similar to previous conclusions (Arbiza et al. 2011; Abadi et al. 2019; Spielman 2019), these results indicate that GTR+I+G should be preferred to existing model selection criteria. However, when branch-lengths estimates were examined, the accuracy measures of GTR+I+G were significantly worse than AIC and BIC for both single-model and complex-model simulation sets (*P < 10*^*-8*^ for all comparisons, Wilcoxon signed rank tests).

### Optimal model for branch-length estimation

We devised ModelTeller, a machine-learning based algorithm optimized to predict the model that yields the most accurate branch-length estimates. To compose the prediction model, we extracted 54 features from each dataset of the abovementioned single-model simulation set (Supplementary Table S2) and used these to train a Random Forest for regression algorithm to predict the ranking of the 24 candidate models in terms of branch-length accuracy. ModelTeller was trained and examined in a 10-fold cross-validation. Namely, the data were divided to ten subsets such that in each iteration, the model was trained on nine subsets and used to make predictions on the 10^th^ subset. This procedure was repeated iteratively until predictions were made for all samples. For performance accuracy, the predicted rankings of models across all datasets were compared to the true rankings (from 1 to 24, being the models that yield the best and worst branch-length estimates, respectively). The average Spearman correlation coefficient between the true and predicted rankings across all datasets was 0.55, and 24% of the datasets resulted in a coefficient above 0.9. In comparison, the Spearman correlation coefficients of AIC and BIC were significantly worse than ModelTeller, such that the average Spearman correlation coefficients between the true rankings and those computed by BIC and AIC were both 0.43 and each resulted in 11% and 13% of the datasets above 0.9 (fig. 1a; *P = 10*^*-138*^ *and 10*^*-145*^ for AIC and BIC, respectively, paired t-test). On average across all datasets, the models that were selected by ModelTeller, BIC, and AIC were ranked 8.19, 8.24, and 8.44 among the true rankings, but these differences were non-significant (fig. 2a, Table 1; *P = 0*.*12* between ModelTeller and AIC, *0*.*41* between BIC and AIC, and *0*.*77* between ModelTeller and BIC, Wilcoxon signed rank test). Altogether, ModelTeller had a marginal improvement over AIC and BIC under the simple scenarios where a single process is assumed to generate the data.

**Table 1.** Average rank of the models selected by the various strategies according to topological and branch-lengths accuracy over the single-model simulations set and the validation set. The table presents the accuracy measures of the trees reconstructed according to the models selected by the various strategies. The reported averages are across all datasets in the single-model simulation set (two left columns) and the validation set (two right columns): (1) Average BS rank: the average rank of the selected model within the ranking of the 24 models according to the BS distances; (2) Average RF rank: the average rank of the selected model within the ranking of the 24 models according to the RF distances (minimal rank for ties without gaps with the increase in rank).

|  | Single-model simulation set |  | Validation set |  |
| --- | --- | --- | --- | --- |
|  | Average BS rank | Average RF rank | Average BS rank | Average RF rank |
| **ModelTeller** | 8.19 | 3.88 | 11.18 | 6.61 |
| **ModelTeller<sub>G</sub>** | 8.16<sup>a</sup> | 3.72 | 11.16<sup>a</sup> | 5.37 |
| **AIC** | 8.44 | 3.51 | 12.61 | 6.57 |
| **BIC** | 8.24 | 3.65 | 12.51 | 6.69 |
| **GTR+I+G** | 10.41 | 3.72 | 13.21 | 5.37 |
| **ModelTest-NG: AIC** | 8.71 | 3.78 | 12.70 | 6.74 |
| **ModelTest-NG: BIC** | 8.43 | 3.84 | 12.60 | 6.82 |
| **AIC<sub>G</sub>** | 8.76<sup>a</sup> | 3.72 | 12.65<sup>a</sup> | 5.37 |
| **BIC<sub>G</sub>** | 8.51<sup>a</sup> | 3.72 | 12.55<sup>a</sup> | 5.37 |
| **Min-BS model<sup>b</sup>** | 1 | 3.30 | 1 | 6.67 |

<sup>a</sup>The ranking of the 24 candidate models was computed where all models were based on a fixed GTR+I+G topology.
<sup>b</sup>The Min-BS model is the model that obtained the minimum BS distance relative to the true tree for each dataset.

**Figure 1.**
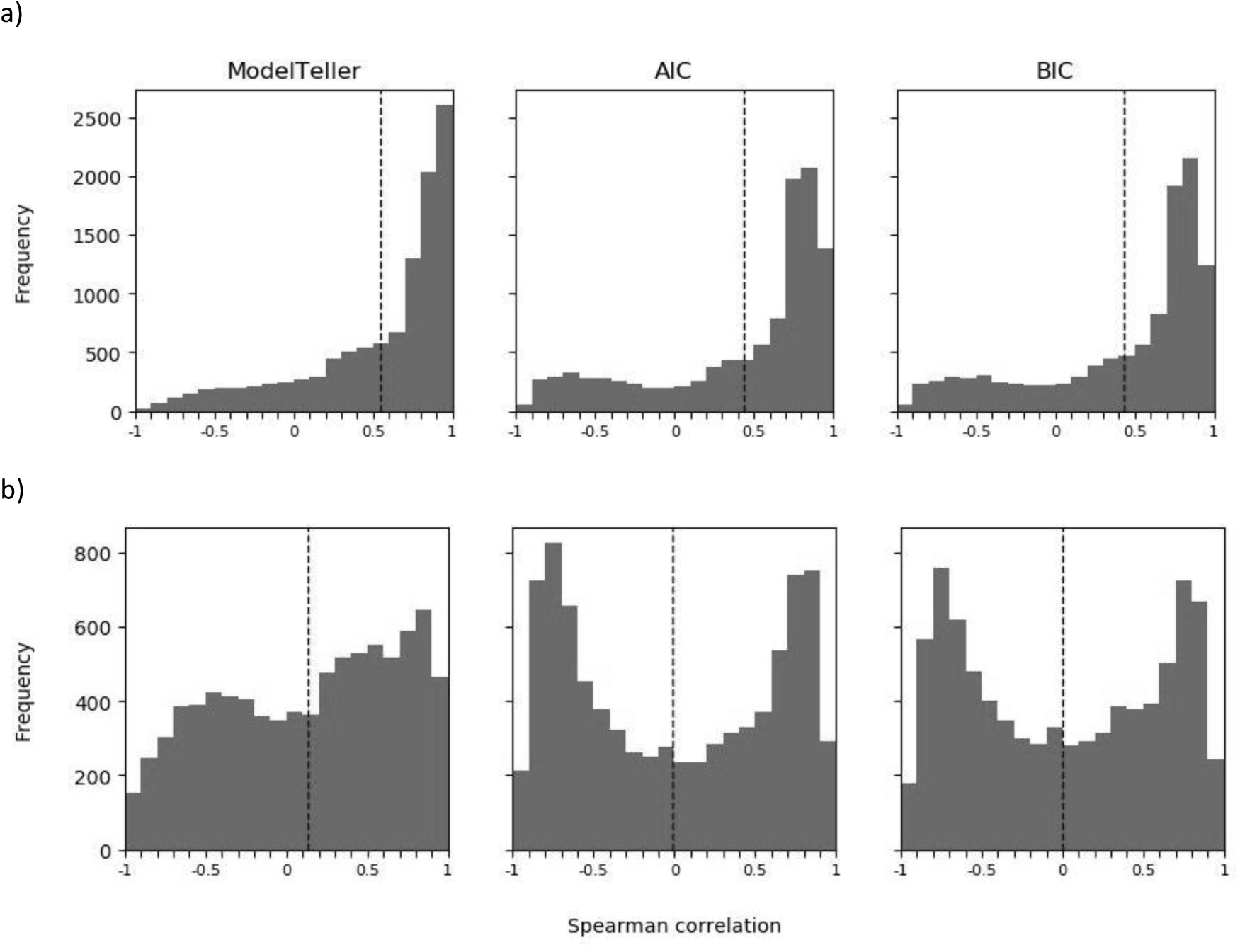
Performance evaluation of ModelTeller and existing model selection criteria. Distribution of Spearman correlation coefficients between the observed and predicted rankings of each of the datasets for (a) the single-model simulation set and (b) the validation set. The x axis represents the Spearman correlation coefficients binned across the interval [-1,1]. The y axis represents the number of datasets for which the correlation between the observed ranking of models and the inferred one corresponds to that coefficient value. The dashed vertical lines represent the averages of each distribution: 0.55 (ModelTeller), 0.43 (AIC), and 0.43 (BIC) for the single model simulations and 0.13 (ModelTeller), −0.02 (AIC), and 0 (BIC) for the validation set.

**Figure 2.**
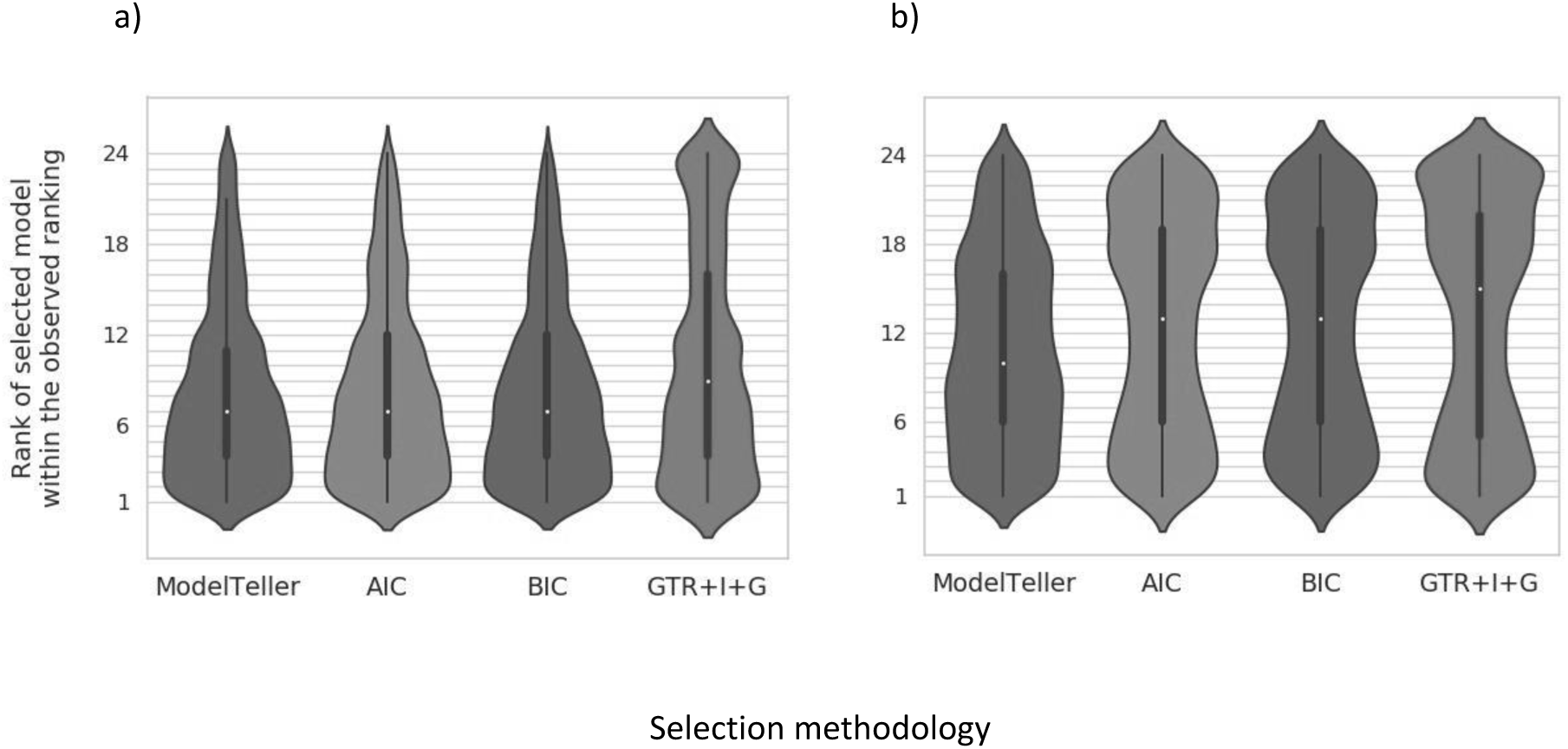
Distribution of the true ranks of the selected models. The x axis represents four methodologies for model selection: ModelTeller, AIC, BIC, and consistently using the GTR+I+G model. The y axis represents the true ranks (from 1 to 24) of the models selected by the methodologies according to the observed BS distance from the true trees. (a) Distribution over the single-model simulation set with medians: 7 for ModelTeller, AIC, and BIC, and 9 for GTR+I+G. (b) Distribution over the validation set with medians: 10 for ModelTeller, 13 for AIC and BIC, and 15 for GTR+I+G. The white dots represent the median, the thick gray bars represent the interquartile (IQR) range, and the thin gray lines extend beyond 1.5*the IQR range. The violin plots represent a kernel density estimation of the underlying distribution of the true rank of the selected models; wider sections of the violin plot represent a higher probability that the methodology selects models of that rank while skinnier sections represent a lower probability.

To examine the performance of ModelTeller on processes that were not used to train it, we simulated a validation set that represents relevant empirical cases. To account for contemporarily relevant cases, the datasets that were employed to estimate the simulations parameters were derived from 8,448 phylogenetic datasets deposited in the heterogeneous TreeBASE public repository (Piel et al. 2009; Vos et al. 2012). To use a simulation procedure different than the one used for training (the single-model simulation set), these datasets were generated using the abovementioned complex-model simulation procedure. ModelTeller, AIC, and BIC were executed for the 8,376 alignments in this validation set. Under these more realistic scenarios, the performances of all strategies were poorer, yet, ModelTeller performed significantly better than AIC and BIC. The models selected by ModelTeller were ranked 11.05 within the true rankings whereas the models selected by BIC and AIC were ranked 12.50 and 12.61 on average (fig. 2b, Table 1; *P = 10*^*-21*^ *and 10*^*-20*^ compared to BIC and AIC, respectively, Wilcoxon signed rank test). The frequencies at which AIC or BIC selected models that produce the top five trees were symmetric to the frequency at which they selected models that produced the worst five trees (median=13; fig. 2b). Exploration of the data showed that AIC and BIC tended toward parameter-rich models (Supplementary Figure S1) and thus generated a distribution pattern that resembles a consistent usage of the most parameter-rich model, GTR+I+G. ModelTeller tended toward models which were ranked better than AIC and BIC (median=10). Similarly, for both BIC and AIC the distribution of the Spearman correlation coefficients was bimodal with increasing frequencies towards the boundaries (+1 and −1) and an average around zero (0.0 for BIC and −0.02 for AIC; fig. 1b), implying that correct ranking of models (from best to worst) was obtained at a similar frequency to ranking the models in a completely reverse manner (from worst to best; fig. 2a). In comparison, the distribution of the Spearman correlation coefficient of ModelTeller presented a pattern by which correct rankings were more frequent than reverse rankings, with a significantly higher average of 0.13 (*P = 10*^*-164*^ *and 1×10*^*-159*^ when ModelTeller was compared to AIC and to BIC, respectively, paired t-test).

### Running time

The rapid feature extraction and their assignment in a readily implemented prediction model grants ModelTeller an advantage over existing model selection criteria that are based on the maximum-likelihood computation, which consist of an iterative optimization of the parameters estimates, topologies, and branch lengths for each candidate model. Over the validation set, the execution of AIC and BIC took 56 minutes on average as opposed to ModelTeller which took 39 seconds on average, an average decrease of a factor of 72 on average.

We also compared the performance of the abovementioned strategies to ModelTest-NG (Darriba et al. 2019), a recent reimplementation of jModelTest that was shown to be faster but not less accurate than the original. Maximum likelihood computations by ModelTest-NG and jModelTest led to similar selections of models for 91% of the datasets for BIC and for 87% of the datasets for AIC. However, branch-lengths accuracy as well as topological accuracy of the inferred phylogenies were inferior compared to AIC and BIC as implemented in jModelTest (and therefore also to ModelTeller), both for the single-model simulations set and for the validation set (Table 1). In spite of the reduced accuracy, the computation by ModelTest-NG took, on average, 3.8 minutes for the validation set, much faster than jModelTest, but still 7 times slower than ModelTeller, on average.

### Optimal model for branch-lengths estimation on a fixed topology

Even though ModelTeller was more accurate in estimating branch lengths compared to AIC and BIC, both in cross validation over the single-model simulation set and over the validation set, it showed lower accuracy when topology reconstruction was examined using the RF distance. For example, among the ranking of the models according to topology reconstruction accuracy, the average rank of the model selected by ModelTeller on the validation set was 6.61 compared to 5.37, 6.56, and 6.69 by GTR+I+G, AIC, and BIC (*P < 10*^*-60*^ between GTR+I+G and each of the others, Wilcoxon signed rank test; see Table 1 for results over the single-model simulation set). Since ModelTeller is optimized at branch-lengths accuracy, we suspected that using the best model for branch-length estimation comes at the cost of poorer topology reconstruction. To examine this hypothesis, for each dataset we detected the best model for branch-lengths estimation, i.e., the model that achieved the highest branch-lengths accuracy (the min-BS model), and measured its rank according to topological accuracy. The average topology rank of the models that were best for branch-length estimates was 6.67, higher (i.e., worse) than its rank by ModelTeller, GTR+I+G, and AIC. This result indicated that when complex patterns underlie the data, reconstruction using one single model does not yield the optimal phylogeny.

In light of this, we devised a two-step phylogeny reconstruction procedure (referred to as ModelTeller_G_), such that a topology is first reconstructed using the GTR+I+G model and then the Random Forest model is trained to predict the ranking of the 24 candidate models in terms of branch-lengths accuracy given the GTR+I+G topology. On the training set, in a 10-fold cross validation, ModelTeller_G_ resulted in an average rank of 8.16 according to branch-lengths. The inferred distances for ModelTeller and ModelTeller_G_ were non-significant (*P = 0*.*08*, Wilcoxon signed rank test). Similarly, on the validation set, models selected for branch-lengths estimation by ModelTeller_G_ were ranked 11.16 according to the branch-length distance. These distances were non-significant when compared to ModelTeller (*P = 0*.*26*) but significantly better than AIC or BIC (*P = 10*^*-35*^ *and 10*^*-34*^ when comparing ModelTeller_G_ to AIC and BIC, respectively, Wilcoxon signed rank test). Despite the similarity to branch-length estimation by ModelTeller, these came with a significant improvement in topology reconstruction, owing to the GTR+I+G topologies.

To examine whether the improvement of ModelTeller over AIC and BIC stems from the better topology reconstruction of the GTR+I+G model, we executed AIC and BIC in a similar two-step manner, i.e., we used a maximum likelihood reconstruction of the GTR+I+G model as a fixed topology, optimized the parameters and the tree branch lengths according to each of the candidate models, and used AIC and BIC for model selection among those (termed AIC_G_ and BIC_G_). When examined over the single-model simulation set, fixing the GTR+I+G topologies did not grant AIC_G_ and BIC_G_ any advantage over AIC and BIC (Table 1). On the validation set, although the GTR+I+G topologies were better than those of AIC and BIC, the average ranks of AIC_G_ and BIC_G_ were marginally higher (i.e., worse) than AIC and BIC, and thus worse than those of ModelTeller_G_ (Table 1).

### Feature importance

The underlying algorithm of ModelTeller, Random Forest, consists of selecting the feature that separates the samples at each split of the tree most accurately during the construction of the decision tree. Eventually, the accumulated effect of a feature across the forest of decision trees reflects its importance for the prediction accuracy. Given a multiple sequence alignment (MSA) of the examined nucleotide sequences, ModelTeller extracts features that could be assigned to three groups (Supplementary Table S2). The first group consists of features that were computed directly from the MSA. These represent the nucleotide composition and measures concerning the similarity between the sequences. For the second group of features, we computed several tree features and estimated the parameters included in the GTR+I+G model using a rapid reconstruction of a BioNJ tree (Gascuel 1997). The third set includes features of the MSA that were computed for a subset of sequences, excluding those of a distant group. This subset was defined by the larger cluster in the partition induced by the longest branch in the BioNJ tree. When the entire set of features is examined, one feature may receive seemingly low importance value due to the presence of a correlated feature (e.g., features indicating the frequencies of different nucleotides within the alignment). To this end, we computed the correlation between the values of every two features and hierarchically clustered them such that the distance of every internal node from the tips defines the minimal correlation between every two features in the induced cluster and internal nodes represent the cumulative importance of the induced cluster (fig. 3). The features with highest importance percentages were the alpha parameter of the gamma distribution and the proportion of invariant sites (21% and 8.7%), which were estimated by an optimization of the GTR+I+G model parameters on the BioNJ tree. Indeed, when ModelTeller selected models that do not include the “+G” component, the median alpha estimation was 20, implying little heterogeneity of rates along the alignment sites in contrast to a median of 1.4 when it was included (*P = 10*^*-170*^ using student t-test; Supplementary Figure S2a). When ModelTeller selected models that include the “+I” component, the median proportion of invariant sites was 0.038, compared to 0.012 for the datasets for which this component was not included (*P = 10*^*-156*^, student t-test; Supplementary Figure S2b). Following these two features, clusters of minimal correlation of 80% with the largest cumulative importance were those associated with sequence divergence attributes (e.g., the mean and variance of the branch-length distribution; cumulative importance of 14.4%), number of alignment sites (e.g., MSA length, number of unique site patterns; cumulative importance of 7.4%), and tree shape attributes (e.g., the average number of branches in a path between every pair of tips; cumulative importance of 6.6%).

**Figure 3.**
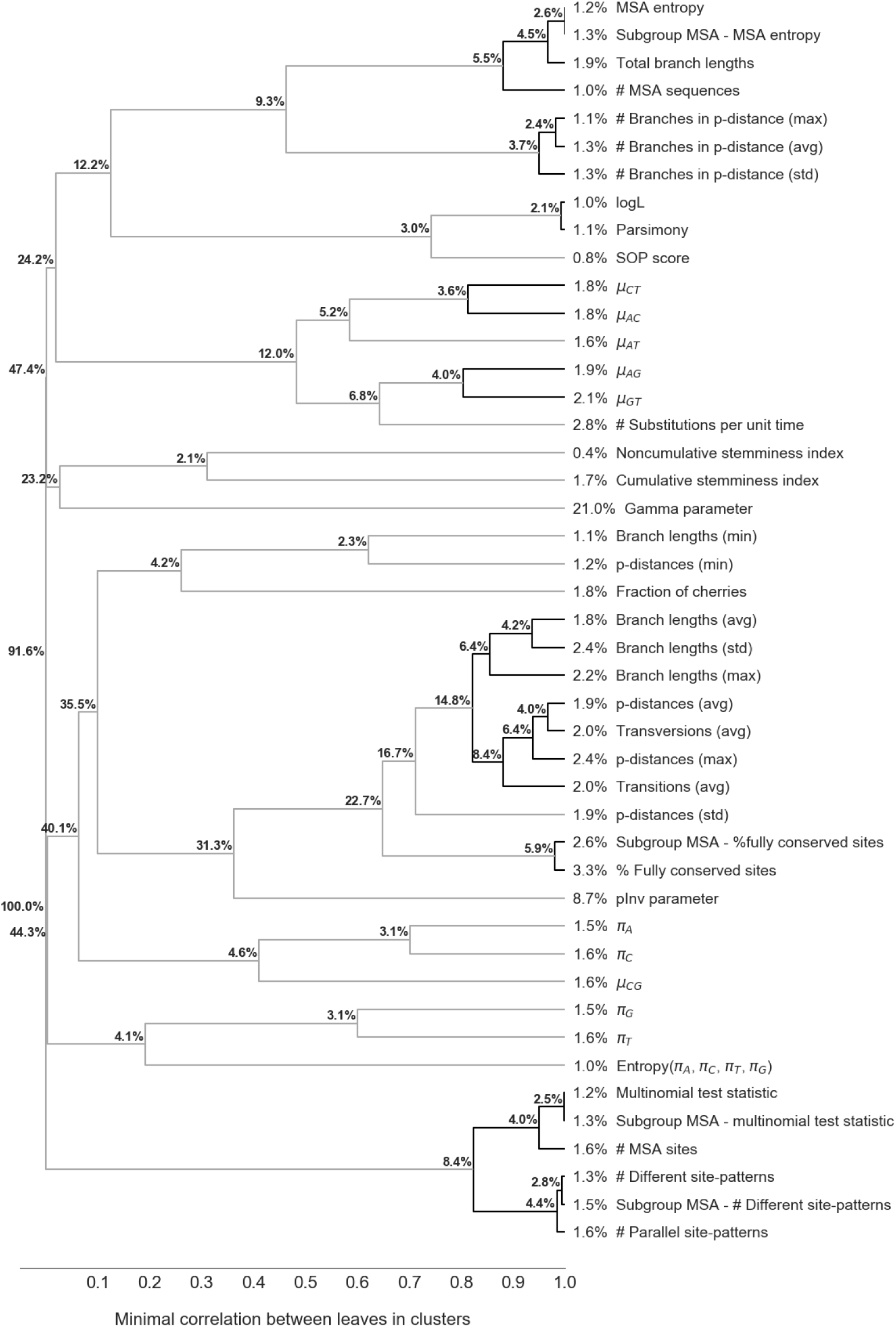
Cumulative importance of clustered features. The dendrogram presents an hierarchical clustering of the features according to the absolute Pearson correlations between them (computed by the complete linkage clustering algorithm, implemented in python SciPy module (Eric et al. 2001)). Node heights represent the minimal correlation between any two features in the induced cluster. Features at the tips of a darker cluster have more than 80% pairwise correlation. The percentages next to the features labels represent their relative importance for prediction. The percentage next to an inner node represents the sum of the importance values of the features that subtend it. Since the models were incorporated as features, their cumulative importance (30%) was excluded from the analysis and the contributions of the presented features were divided by their sum (for the actual importance percentages and the description of the features, see Supplementary Table S2).

## Discussion

For decades, phylogenetic model selection has indisputably relied on likelihood computations of all alternative models. Many statistical criteria have been proposed over the years, focusing on the inference task of understanding the processes that underlie the data at hand. In recent years, various learning tools have emerged, aimed at learning the patterns within the data for identifying the best course of action for a certain prediction task. In this study, we utilized the machine-learning framework to directly predict the optimal model for our desired prediction task – the reconstruction of the phylogenetic tree for a given set of sequences. Because the tree topology is quite robust to the model choice (Buckley et al. 2001; Posada and Crandall 2001; Sullivan and Swofford 2001; Abdo et al. 2005; Sullivan et al. 2005; Kelchner and Thomas 2007; Ripplinger and Sullivan 2008; Hoff et al. 2016), we have focused here on the accuracy of the estimated branch lengths.

Notably, two major goals in the study of biological systems are inference and prediction. In phylogenetics, statistical inference is an important component to identify the processes that generated the data, but carries multiple drawbacks. First, not all statistical criteria have been adjusted to phylogenomics. For example, AIC and BIC assume independence of the data particles, an assumption that does not hold for sequence data. Second, statistical criteria rely on an extensive search throughout the parameter space and optimization of the likelihood score or its components. These searches entail long computation time. The running time also increases considerably with the size of the data. The abundance of parameters and the accumulating amounts of data often make the computation infeasible, mainly for Bayesian methods. Third, the goal that is achieved by applying these methods is identifying the underlying processes rather than the ultimate goal: the accuracy of the reconstructed phylogeny. As shown in our study, the existing maximum likelihood criteria perform very well at revealing the underlying process, at least when it is relatively simple, but these models do not necessarily lead to the most accurate phylogenetic tree. In contrast, the learning paradigm makes minimal assumptions on the data generating process and is aimed at predicting the desired outcome. Once a trained model is set up, the prediction process is rapid and the running time relies merely on the extraction of informative features that describe the data.

With the accumulation of massive biological databases and advancements in automating data extraction, many biological frameworks have converted from inference of the underlying processes to generalizing the links between the patterns and the outcome in the observed data for prediction of future samples. For example, the amplification of brain measurements has shifted the research of cognitive neurosciences from isolating the effect of one variable to learning complex brain patterns, thus enabling prediction of future brain scans (Norman et al. 2006; Bzdok 2017). Studying disease profiles based on microarray data has advanced from attempting to detect a single gene that induces a condition in correlation tests to learning the complex interactions between genes and to predicting the condition (Guyon et al. 2002). Notwithstanding, machine-learning poses several limitations that should be addressed while setting up the training data. First, models that were not included in the training of ModelTeller cannot be assessed in the prediction process. Here, we used the nucleotide substitution models that are most commonly used as candidate models in phylogenetic model selection applications (Stamatakis et al. 2005; Posada 2008; Darriba et al. 2012). However, this is only a subset of all possible substitution models, and when new models are proposed the learning machinery should be retrained. Second, machine learning cannot make accurate predictions for patterns that were not found in the training set. To minimize this limitation, we assembled a large cohort of empirical data that extends over a large range of realistic patterns. We also showed that when given data that were generated from evolutionary patterns that are different from the ones it was trained on, ModelTeller outperformed AIC and BIC.

The data that was used to train ModelTeller presents low level of complexity and does not account for possible model violations because they were generated according to a single model which is also part of the set of candidate models. ModelTeller did not overwhelmingly outperform AIC and BIC on this training set. However, when examined over the validation set, in which data were drawn from a public repository of contemporary investigated cases and the simulation patterns were more intricate than a single simplistic model, ModelTeller selected better models and also ranked the candidate models more accurately (fig. 2). For this set, the rankings of AIC and BIC presented a symmetric pattern whereby successful rankings (with a Spearman correlation coefficient approaching 1) were as frequent as totally inverse ones (with a Spearman correlation coefficient approaching −1; fig. 1a). Furthermore, the models that were selected by AIC and BIC were the best or the worst ones at similar frequencies (fig. 2a). Examination of the selections showed that AIC and BIC tended to select complex models for these datasets, thereby presenting a pattern that is similar to consistently selecting the most complex model, GTR+I+G, which was the best model and worst model at similar frequencies. These selections perhaps better describe the generating process, which was more complex than any other alternative, but do not lead to more accurate branch-length estimates.

Ultimately, empirical sequence data would have been the best choice for a training set. Evidently, the true phylogenies of empirical datasets are unknown. An alternative would be to train the machine-learning on data that are simulated according to intricate cases that mimic realistic processes, similar to the validation set used in this study. Other realistic simulation scenarios were previously proposed. Philippe et al. (Philippe et al. 2005) performed simulations based on combinations of within-site rate variation (heterotachy) and variation of evolutionary rates across lineages to show that maximum-likelihood methods are more accurate than maximum-parsimony. Sipos et al. (Sipos et al. 2011) devised a simulator that uses the Gillespie algorithm to integrate the actions of many concurrent processes such as substitutions, insertions, and deletions rather than assuming a homogeneous substitution model. Nevertheless, no matter which simulation method is employed, it must extend over the large range of possible realistic processes so that the trained model is applicable for many empirical datasets.

Even though ModelTeller was at least as accurate as AIC and BIC, it still did not reveal the optimal model with high frequency. Within a ranking of 1 to 24, the average rank of the models predicted by ModelTeller was 8.19 on the training data and 11.18 on the validation data. Obviously, recording the rank of the selected models does not reflect the actual distances of the reconstructed trees, therefore this measure cannot indicate whether one model leads to substantially different inferences than another. However, since the tree sizes in the data vary, measuring the ranks of models enables to distinguish between better and worse selections regardless of the tree size. Due to these reasons, we incorporated the ranking in the learning phase, such that the prediction model was trained to predict the ranks of the different candidate models. In a simple classification task, the machine-learning algorithm could have been trained to identify the single most accurate model. While in trials this procedure predicted the best model for 4% additional datasets (data not shown), when it selected an inaccurate model, this model was of a very low rank. One advantage of predicting the ranking of models is that it not only enabled learning which models are preferred but also which are unfavorable per dataset. Furthermore, in case that only a subset of the candidate models is implemented in downstream application software, the user can use the first model that is available among the predicted ranking.

The learning procedure allowed us to measure the contribution of features that have previously been inspected with respect to determining a substitution model for tree reconstruction. To this end, we included the alignment size, which is used in the penalty function of BIC and AICc, and several test statistics that were proposed for testing model adequacy, such as the percentage of fully conserved sites, the distinct site-patterns, and parallel site-patterns suggested by Goldman (1993b), and the multinomial test statistic suggested by Bollback (2002). All of these features resulted in similar contribution to the prediction model in the feature importance analysis (between 1.2% to 3.3%). Notably, the effect of each of these features was obscured by other features. For example, the number of parallel site-patterns, i.e., sites that correspond to a similar pattern of evolution regardless of the identity of the nucleotide (e.g., ACCCAA and AGGGAA correspond to the pattern XYYYXX) were highly correlated with the number of distinct site-patterns (in which ACCCAA and AGGGAA are counted as two patterns; Pearson *r*^*2*^ *= 0*.*99*), and together, these features obtained an importance of 4.4%. Perhaps surprisingly, the importance of these correlated features was higher than that of the two features that represent the alignment size most intuitively; the importance of the number of sequences was only 1% while that of the alignment length was 1.6%. This suggests that computing the information criteria for phylogenetic model selection might be improved by addressing the data size as the number of independent sites instead of the prevailing approach for addressing the data size as the number of alignment sites.

A benefit of the machine-learning approach for model selection is the possibility to change the target function for any desired inferential task. In this study we used the Branch-Score (BS) distance proposed by Kuhner and Felsenstein (Kuhner and Felsenstein 1994), a distance between unrooted trees that is most sensitive to the accuracy of branch-length estimates. However, different distances could be employed as well. For example, the Robinson-Foulds (RF) distance (Robinson and Foulds 1981) could have been employed if topological accuracy had been the focus. This distance counts the number of branch partitions that appear in one tree but not the other, scoring 1 for each unmatched partition. Since it was previously shown that the inferred topology is robust to the selection of model, performing model selection for this goal is not expected to be beneficial (Buckley et al. 2001; Posada and Crandall 2001; Sullivan and Swofford 2001; Abdo et al. 2005; Sullivan et al. 2005; Kelchner and Thomas 2007; Ripplinger and Sullivan 2008; Arbiza et al. 2011; Hoff et al. 2016; Abadi et al. 2019). Kuhner and Yamato (Kuhner and Yamato 2015) examined the performance of nine tree comparison methods, and concluded that branch-length based distances are more suitable for comparison of similar trees (as are the true tree and inferred one) than those based on topological difference only, even if topology is the focus. Among those, they found that a length based variation of the Robinson-Foulds (RFL) distance (Robinson and Foulds 1979) is the most accurate for rooted trees, which is quite similar to the BS distance we used. Both RFL and BS increment the distance by the difference between corresponding branches, however, the absolute value of this difference is used in the former and the squared value in the latter. Using the BS distance grants higher weights to large differences between branches and lower weights to small differences, but since we compared trees inferred from the same data but with different models, we do not expect this subtle difference to change the results to a large extent. In the abovementioned study, Kuhner and Yamato studied additional metrics but most of them are restricted to clocklike rooted trees, which puts additional constraints on the data the prediction model is applicable to. Other restrictions on the data and the utility could be to analyze only coding or non-coding sequences, thus refining the accuracy of the learning procedure for specific niches. A similar prediction model could be trained over models of codons or proteins or a combination of different models for partitions of the sequence data. For such variations, the current framework of ModelTeller could serve as a foundation, but new data must be generated, relevant features should be extracted, and a new learning model should be trained.

## Methods

### Data assembly

The data used in this study for learning were simulated using characteristics and rates derived from empirical alignments. The alignments were obtained from four databases that differ in the evolutionary relationships between the sequences: (1) OrthoMam (Ranwez et al. 2007; Douzery et al. 2014), a database of orthologous mammalian markers; (2) Selectome (Moretti et al. 2014), which includes codon alignments of species within one of four groups (Euteleostomi, Drosophila, Primates, and Glires; to avoid overlap with OrthoMam, the last two groups were excluded); (3) PANDIT (Whelan et al. 2003), which includes alignments of the nucleotide coding sequences of protein domains; (4) ProtDB(Carroll et al. 2007), which includes genomic sequences that were aligned according to the tertiary structure alignments of the encoded proteins published in BALIBASE (Thompson et al. 2005), SMART (Ponting et al. 1999), OXBench (Raghava et al. 2003), and Prefab (Edgar 2004). From these four databases, alignments were sampled among those with 5 to 500 taxa and 50 to 5,000 alignment sites. From OrthoMam, Selectome, and PANDIT, 3,000 alignments were sampled to represent a broad spectrum of data size. Specifically, each database was divided to 20 bins according to the number of sequences, and 150 alignments were randomly sampled from each bin. From ProtDB, 1,270 datasets that comply with this range of data size were included. Together, these data included 10,270 datasets that range in sequence divergence, number of taxa, and alignment length. In a preliminary examination, we observed that a large fraction of the sampled datasets contain sequences of low divergence but only few with high divergence (reflected by the summation of branch lengths in the reconstructed trees). Therefore, we included additional datasets whose sequence divergence is large. To this end, for each dataset that was not sampled from the databases we computed the BioNJ tree using PhyML and added those for which the summation of branch lengths was greater than ten, resulting in 563 additional datasets. Altogether, the studied set contained 10,832 datasets.

#### Single-model simulation set

For each dataset, we performed simulations according to one of 24 models whose rates were derived from the empirical dataset. To this end, PhyML was executed for each empirical dataset using a randomly selected model to infer the phylogenetic tree (established as the true tree) and the model parameters. The random selection was among a set of 24 substitution models, i.e., JC, F81, K2P, HKY, SYM, and GTR – each one with the presence/absence of the +I (proportion of invariant sites) and +G (heterogeneity of rates among sites following the discrete gamma distribution with four categories) options. Given the selected model, the inferred tree, and the model parameters, an alignment was simulated using INDELible (Fletcher and Yang 2009).

#### Complex-model simulation set

To enhance the complexity of the simulations, we simulated additional datasets with heterogeneity of models and rates across the alignment sites rather than simulating according to a single substitution model for the entire alignment. For the first layer of complexity, variation in the substitution pattern across the alignment sites, each empirical dataset was divided to partitions of 50 sites (datasets were trimmed such that the alignment length is divisible by 50). jModelTest (Guindon and Gascuel 2003; Darriba et al. 2012) was executed for each partition to obtain the best-fitted model and its inferred free parameters. To obtain the best-fitted model while avoiding a bias toward a particular criterion, one maximum-likelihood criterion, i.e., AIC or BIC, was randomly selected per partition. The second layer of complexity was obtained by providing the simulator site-specific rates that were drawn from an empirical distribution, fitted to each dataset by Rate4site (Mayrose et al. 2004), rather than being drawn from the gamma + invariance (G+I) distribution. To combine these two layers of complexity, INDELible (Fletcher and Yang 2009) was used to simulate each site given its respective model and rate and the simulated sites were concatenated to form a single alignment. The input trees in these simulations were reconstructed from the respective empirical alignments using BioNJ (Gascuel 1997) (as implemented in PhyML) with the distance matrix computed using the JC model. These trees were regarded as the true trees for the relevant comparisons. 16 datasets for which jModelTest or Rate4site failed were excluded from this analysis.

### Tree inference, distance metrics, and ranking of models

To determine which model is best for phylogeny reconstruction for each simulated dataset, the maximum-likelihood tree of each of the 24 candidate models was reconstructed using PhyML (Guindon et al. 2010), their distances from the true tree were measured, and the model that yielded the minimal distance was considered as best. For branch-length estimation accuracy, the Branch Score (BS) distance (Kuhner and Felsenstein 1994) was computed, i.e., sum of the squared differences between corresponding branches of the trees, as implemented in Treedist (Felsenstein 2008). Branches that were not found in one tree due to different topologies were treated as having a length of 0. For topological accuracy, the Robinson-Foulds (RF) distance (Robinson and Foulds 1981) as implemented in TreeCmp (Bogdanowicz et al. 2012) was computed. The ranking of models were determined from lowest to highest distances (ranks 1 to 24). Ties were assigned the same rank with no skips between increasing ranks. Since different models almost conclusively lead to different branch-lengths values, the branch-lengths rankings resulted in 24 distinct ranks. In contrast, different models may lead to identical topologies, and so the topologies ranking resulted in fewer distinct ranks. For analysis of a fixed GTR+I+G topology (i.e., ModelTeller_G_, AIC_G_, BIC_G_) the maximum-likelihood phylogeny was first reconstructed in PhyML using the GTR+I+G model, and then the 24 models were optimized along with the branch-length estimates given the GTR+I+G topology. The BS distances of the resulting trees from the true trees were measured. For analysis of the branch lengths given the topologies of the true trees, the 24 models were optimized along with the branch-length estimates given the true trees.

### Machine-learning training and cross-validation

ModelTeller was implemented as a Regression task where the objective is to learn the ranking of the candidate models according to the BS distance between the true and inferred trees. To incorporate all distances within the training phase, each dataset, represented by the extracted features, was replicated 24 times corresponding to the 24 models. Each replicate was assigned with additional features that indicate the respective model (see the ‘Predictive features’ section). The target value for each replicate was the rank of the corresponding model from 1 to 24 (the models that produce minimal and maximal BS distance, respectively; termed ‘the true ranking’). Then, the Distributed Random Forest algorithm implemented in the H_2_O platform (H2O.ai Team 2015) was trained over this extended training set of 10,832 × 24 samples using 50 decision trees. The predicted ranks of the test set were determined by applying the trained algorithm to the test samples replicates and ranking the predicted values from low to high (1 to 24, being best to worst; termed ‘the predicted ranking’). For the cross-validation procedure, in each iteration one decile of the datasets, including all 24 replicates of each, was reserved as a test set, while the others were used for training. In each such iteration the algorithm was trained over the respective training set and the test set was used for prediction of the ranks. We used two metrics for performance evaluation. First, for each dataset we computed the Spearman correlation coefficient between the true ranking of models and the ranking predicted in cross validation (or the one produced by AIC or BIC) and averaged the coefficients across all. Additionally, we averaged the true ranking of the model that was ranked first by the model selection strategy.

### Predictive features

Each dataset was represented by a set of 54 explanatory attributes extracted from it. These features can be classified into three sets of features. The first set are those extracted solely from the sequence alignment. These include the number of sequences, the length of the alignment, the frequency of each type of nucleotide, the alignment entropy, the multinomial statistic, and the sum-of-pairs score (all features are detailed in Supplementary Table S2). The second set of features represent an approximation of the model parameters and the reconstructed maximum-likelihood tree. That is, a basic BioNJ (Gascuel 1997) tree was reconstructed using PhyML (Guindon et al. 2010) and the rate parameters of the GTR+I+G model were optimized on a fixed topology and branch lengths to avoid long computation time (for ModelTeller_G_, those features were computed over the optimized GTR+I+G phylogeny that was computed as a first step). The values of the estimated parameters including the substitution rates, the shape parameter of the gamma distribution (heterogeneity across sites), and the proportion of invariant sites, as well as the sum of branch lengths of the reconstructed phylogeny were used as features. The third set of features describe the similarity within a subset of the sequences, considering that a distant group may affect the prediction accuracy. To select these sequences, the BioNJ tree was pruned at the longest branch, and the sequences of the larger subtree were used to extract the induced MSA from the original one. Then, some of the MSA features (set 1) were computed over this reduced MSA. For a full description of the features and their extraction procedures, see Supplementary Table S2.

The 24 candidate models were represented by four categorical features: (1) the number of parameters in the substitution matrix: 1 for JC and F81, 2 for K2P and HKY, or 6 for SYM and GTR; (2) whether to allow for unequal base frequencies, i.e., 0 for JC, K2P, and SYM or 1 for F81, HKY; GTR (3) whether it accounts for the proportion of invariable sites, i.e., 1 for the inclusion of the +I component and 0 otherwise; (4) whether it accounts for heterogeneous rates across sites, i.e., 1 for the inclusion of the +G component and 0 otherwise.

### Validation set

As a validation set, we used data and simulation procedures different from those used for training ModelTeller. To this end, the data characteristics for the validation set were obtained from TreeBASE (Piel et al. 2009), a repository of user-submitted phylogenies. All available nucleotide alignments were downloaded from the repository. Datasets with fewer than 50 alignment sites or invalid nucleotide alignment characters were excluded. The final validation set comprised of 8,448 datasets. The data characteristics were extracted from the datasets and used to simulate new alignments using the complex-model simulation procedure.

### Implementation and availability

ModelTeller was implemented in python using the Distributed Random Forest algorithm, implemented in the python H2O platform (H2O.ai Team 2015). The score provided by the algorithm, essentially represents the ranking of the models, such that the first one is predicted to yield the best branch-lengths estimation. The ModelTeller utility is available for online use and for download at http://ModelTeller.tau.ac.il/ and offers three running modes: (1) predicting a single model for optimization of the model parameters, the branch lengths, and the topology (ModelTeller); (2) predicting a model for branch-lengths optimization over a fixed GTR+I+G topology (ModelTeller_G_); (3) predicting a model for branch-lengths optimization over a fixed topology given by the user. The online tool provides the predicted ranking of models as well as the output phylogeny for the best model, computed by PhyML (Guindon et al. 2010).

## Data availability

The datasets contained within the single-model, complex-model, and validation simulation sets have been deposited in Open Source Framework (OSF) with the identifier DOI 10.17605/OSF.IO/2WS3A.

## Acknowledgements

S.A. is supported by PhD fellowships provided by the Rothschild Caesarea Foundation and the Edmond J. Safra Center for Bioinformatics at Tel-Aviv University, and travel fellowships provided by Joan and Jaime Constantiner Institute for Molecular Genetics and the Society of Systematic Biology. O.A. is supported by PhD fellowships provided by the Edmond J. Safra Center for Bioinformatics at Tel-Aviv University, the Dalia and Eli Hurvits Foundation LTD, and the Ministry of Science, Technology & Space of Israel. T.P. is supported by an Israeli Science Foundation number 802/16. I.M. is supported by an Israeli Science Foundation number 961/17.

