## Supplementary information for "ModelTeller: model selection for optimal phylogenetic reconstruction using machine learning"

| **Contents** |  |
| --- | --- |
| **S1 Figure:** Distribution of selection of models by the strategies on the validation set | 2 |
| **S2 Figure:** Effect of the pInv and alpha parameters estimates on the selection of ModelTeller | 3 |
| **S1 Table.** Complete features set | 4 |
| **S2 Table.** Complete features set | 5-6 |
| **Supplementary References** | 7 |

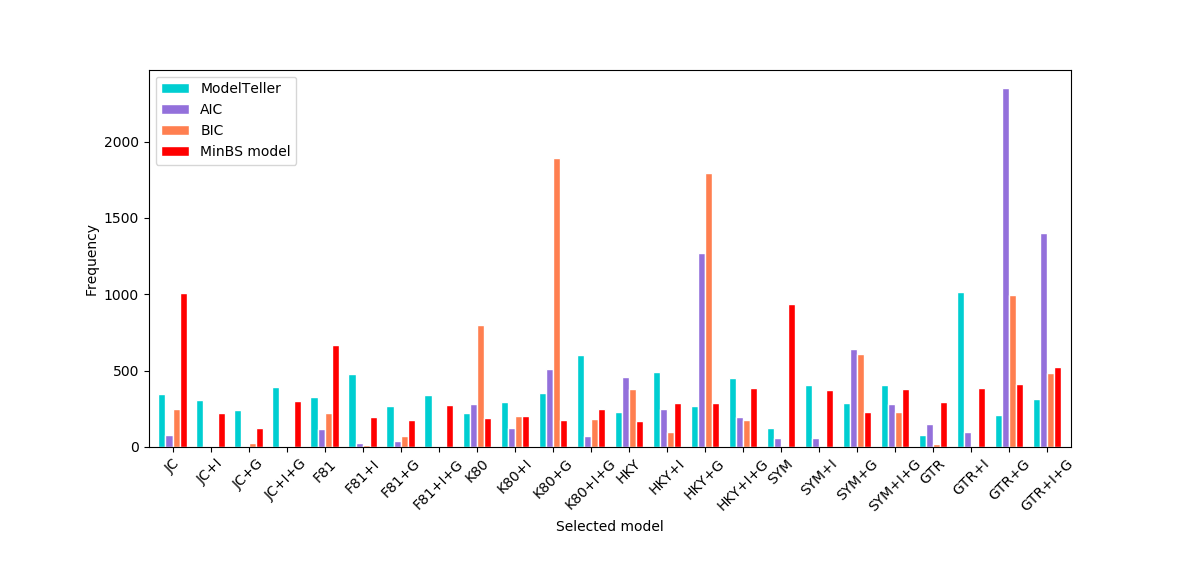

**Figure S1. The Distribution of models selected by the examined strategies on the validation set.** The figure represents the frequency (y-axis) at which each model (x-axis) was selected by each strategy. For each of the 24 models at the x-axis, the different model selection strategies are colored according to the legend at the top left corner and are ordered from left to right: ModelTeller, AIC, BIC, and the minBS model (the model with the lowest BS distance in each dataset).

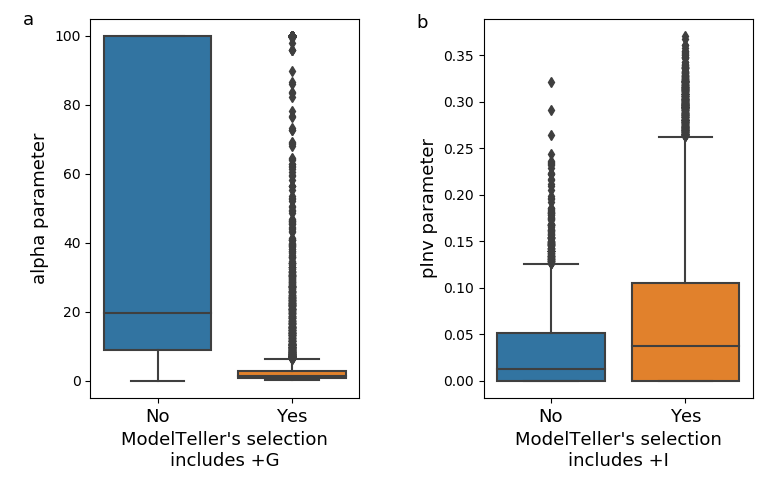

**Figure S2. Effect of the pInv and alpha parameters estimates on the selection of ModelTeller.** (a) The distributions of the alpha parameter (shape of the Gamma distribution for determining rates heterogeneity across the alignment sites) for the models selected by ModelTeller. The blue and orange (left and right) boxplots represent models with or without the “+G” component in the selected model (significantly different, *P = 10^-170^*, student t-test). (b) The distributions of the pInv parameter (proportion of invariant sites) for the models selected by ModelTeller. The blue and orange (left and right) boxplots represent models with and without the “+I” component in the selected model (significantly different, *P = 10^-156^*, student t-test). The boxes extend from the lower to upper quartile values of the data, with a line at the median. The whiskers reach 1.5 times past the first and third quartiles. Flier points are the values past the end of the whiskers.

|  | **Single-model simulations** | | | | **Complex-model simulations** | | |
| --- | --- | --- | --- | --- | --- | --- | --- |
| **Strategy** | | **RF^1^**  **avg (std)** | **Ranked RF^2^**  **avg (std)** | **% identical^3^** | **RF^1^**  **avg (std)** | **Ranked RF^2^**  **avg (std)** | **% identical^3^** |
| **AIC** | | 2.51 (6.92) | 1.99 (0.27) | 42% | 2.94 (7.50) | 2.01 (0.29) | 38% |
| **BIC** | | 2.54 (6.98) | 2.00 (0.31) | 42% | 2.97 (7.54) | 2.03 (0.35) | 38% |
| **GTR+I+G** | | 2.57 (7.18) | 2.00 (0.46) | 42% | 2.90 (7.48) | 1.96 (0.45) | 39% |
| **p-value** | | AIC ↔ GTR+I+G: 0.044  AIC ↔ BIC: 0.013  BIC ↔ GTR+I+G: 0.8 | | | AIC ↔ GTR+I+G: 3x10^-7^  AIC ↔ BIC: 7x10^-4^  BIC ↔ GTR+I+G: 1x10^-15^ | | |

|  | **Single-model simulations** | | **Complex-model simulations** | |
| --- | --- | --- | --- | --- |
| **Strategy** | **Bs^4^**  **avg (std)** | **Ranked Bs^5^**  **avg (std)** | **Bs^4^**  **avg (std)** | **Ranked Bs^5^**  **avg (std)** |
| **AIC** | 0.285 (0.48) | 1.89 (0.64) | 0.282 (0.32) | 1.98 (0.60) |
| **BIC** | 0.283 (0.47) | 1.86 (0.54) | 0.281 (0.32) | 1.96 (0.80) |
| **GTR+I+G** | 0.294 (0.51) | 2.25 (0.91) | 0.282 (0.31) | 2.05 (0.83) |
| **p-value** | AIC ↔ GTR+I+G: 1x10^-207^  AIC ↔ BIC: 3x10^-5^  BIC ↔ GTR+I+G: 2x10^-192^ | | AIC ↔ GTR+I+G: 2x10^-8^  AIC ↔ BIC: 3x10^-4^  BIC ↔ GTR+I+G: 7x10^-12^ | |

**Supplementary Table S1. Comparison of topology reconstruction and branch-length estimation between AIC, BIC, and consistently use of the GTR+I+G model.** The tables present the accuracy measures of the trees reconstructed according to the models selected by AIC, those selected by BIC, and GTR+I+G compared to the true trees that were used to simulate the data. The single-model simulations set refers to the training set, i.e., each alignment was simulated according to a single model and its parameters estimated on an empirical dataset. The complex-models simulations set was generated according to parameters estimated on the same data, but assuming heterogeneity of models and rates across the alignment sites. The reported averages, standard deviations (in parenthesis), and percentage are across all datasets. The top table presents the topological accuracy and the bottom table presents the branch-length estimation accuracy. (1) RF: the Robinson-Foulds (Robinson and Foulds 1981) distances between the inferred topologies and those of the true trees; (2) Ranked RF: for each dataset, the RF distances of the three strategies were ranked from one to three (average rank for ties); (3) %identical: the percentage of inferred topologies that were identical to the true trees; (4) Bs: the Branch-Score (Felsenstein 2008) distances, i.e., the Euclidean distance between the branch lengths of the inferred phylogenies and those of the true trees; (5) Ranked Bs: for each dataset, the Bs distances of the three strategies were ranked from one to three (average rank for ties). The p-values are for a Wilcoxon signed-rank test between the distances generated by every two strategies (since the test is nonparametric, these values are identical for the absolute distances and the ranked ones). P-values marked in red are considered significant following a Bonferroni correction for multiple testing, below a threshold of 0.05.

| **Feature importance** | **Label** | **Description and computation method** |
| --- | --- | --- |
| **Model components**  The model was integrated as a combination of four categorical features.  These features and their contribution were excluded from the analysis presented in Fig. 3. | | |
| 2.13% | F component | Whether to allow for unequal base frequencies, i.e., 0 for JC, K2P, and SYM or 1 for F81, HKY, and GTR |
| 15.95% | G component | Whether it accounts for heterogeneous rates across sites, i.e., 1 for the inclusion of the +G component and 0 otherwise |
| 7.28% | I component | Whether it accounts for the proportion of invariable sites, i.e., 1 for the inclusion of the +I component and 0 otherwise |
| 4.66% | Matrix component | The number of free substitution parameters: 1 for JC and F81, 2 for K2P and HKY, or 6 for SYM and GTR |
| **Set 1: MSA (Multiple Sequence Alignment) features**  The features below were computed over the MSA using Python scripts | | |
| 1.25%, 1.32%, 1.13%, 1.10%, 1.25%, 1.46% | $\mu_{AC}$,$\mu_{AG}$,$\mu_{AT}$,$\mu_{CG}$,$\mu_{CT}$,$\mu_{GT}$ | Nucleotide-to-nucleotide subsitutions between every pair of sequences averaged across all pairs in the MSA |
| 1.43% | Transitions (avg) | Average of $\mu_{AG}$ and $\mu_{CT}$ |
| 1.42% | Transversions (avg) | Average of $\mu_{AC}$, $\mu_{AT}$, $\mu_{CG}$, and $\mu_{GT}$ |
| 1.03%, 1.11%, 1.06%, 1.10% | $\pi_{A}$, $\pi_{C}$, $\pi_{G}$, $\pi_{T}$ | The frequency of A, C, G, and T nucleotides observed in the MSA |
| 0.72% | Entropy($\pi_{A}$, $\pi_{C}$, $\pi_{G}$, $\pi_{T}$) | Shannon entropy (Shannon 1948) computed over $\pi_{A}$, $\pi_{C}$, $\pi_{G}$, and $\pi_{T}$ |
| 0.71% | MSA entropy | Shannon entropy (Shannon 1948) over the alignment sites |
| 0.83% | MSA multinomial test statistic (Bollback 2002) | MSA multinomial test statistic (Bollback 2002)  $T\left( X \right)=\left( \sum_{i=1}^{n} N_{\xi\left( i \right)}\ln(N_{\xi\left( i \right)}) \right)-N \ln(N)$  $N_{\xi\left( i \right)}$ – the number of times the unique pattern $\xi$ of site $i$ is observed, among $n$ unique patterns |
| 0.90% | # Different site-patterns (Goldman 1993) | Number of different MSA columns and their fraction out of the MSA length |
| 1.11% | # Parallel site-patterns (Goldman 1993) (*) | Number of parallel columns, e.g., the columns ACCCAA, CAAACC, AGGGAA are parallel and correspond to the pattern XYYYXX(Goldman 1993) |
| 1.10% | # MSA sites | Number of positions in MSA (MSA length) |
| 0.72% | # MSA sequences | Number of sequences in MSA |
| 2.33% | % Fully conserved sites (Goldman 1993) | Percentage of fully conserved sites in MSA |
| 0.57% | SOP score | Sum-of-pairs score with score of 1 for a match, -1 for a mismatch and a gap. Scores were summed for every pair of sequences and across all pairs in MSA |
| **Set 2: Initial tree features**  PhyML(Guindon et al. 2010) was executed for the MSA to reconstruct a basic BioNJ tree with optimization of the GTR+I+G model parameters only, i.e., the tree topology and branch lengths were not optimized.  The features below were either extracted from the output of PhyML or computed over the reconstructed tree. | | |
| 0.78% | Parsimony | The number of nucleotide substitutions along the tree |
| 0.72% | logL | log Likelihood |
| 14.71% | Gamma parameter | The shape parameter of the gamma distribution (heterogeneity across sites) |
| 6.06% | pInv parameter | Proportion of invariant sites |
| 1.97% | # Substitutions per unit time | Extracted from the rate matrix in PhyML output. This rate, $r$, is computed by dividing the instantaneous substitution rate of nucleotide $X$ to $Y$, i.e., $r\mu_{XY}\pi_{Y}$ (an entry in the matrix), by the substitution rate $\mu_{XY}$ and $Y$ stationary base frequency $\pi_{Y}$ |
| 1.35% | Total branch lengths | Sum of tree branch lengths |
| 1.54%, 0.80%, 1.25%, 1.67% | Branch lengths (max, min, avg, std) | Maximum, minimum, average, standard-deviation, and Shannon entropy (Shannon 1948) were computed over the set of tree branches |
| 1.69%, 0.83%, 1.36%, 1.67% | p-distance (max, min, avg, std) | p-distance – the sum of branch-lengths in a path between two tips. Maximum, minimum, average, standard-deviation, and Shannon entropy (Shannon 1948) were computed over the set of all tree p-distances |
| 0.80%, 0.89%, 0.93% | # Branches in a p-distance (max,  avg, std) | The number of branches in a path between two tips. Maximum, average, standard-deviation, and Shannon entropy (Shannon 1948) were computed over the set of the branch number in all tree p-distance |
| 1.20% | Cumulative stemminess index (Fiala and Sokal 1985; Ripplinger and Sullivan 2010) | Stemminess index of a node is computed as the ratio between the length of the branch (“the stem”) and the size of the subtree induced by the node (the sum of branches together with the stem for the cumulative (Fiala and Sokal 1985) index, and the maximal distance from the tips for the noncumulative index (Rohlf et al. 1990)). The stemminess index of the tree is the average stemminess indices of all internal nodes excluding the root. To root the tree, an ‘outgroup’ was set according to the longest branch |
| 0.31% | Noncumulative stemminess index (Rohlf et al. 1990; Mooers 2004) (*) |  |
| 1.29% | Fraction of cherries (McKenzie and Steel 2000) | Clades that consist of exactly two tips divided by the total potential number (1/2 of the number of tips) |
| **Set 3: Subgroup MSA features**  Two subgroups were determined according to the partition induced by the longest branch in the non-optimized GTR+I+G tree described above. | | |
| 0.90% | Subgroup MSA - multinomial test statistic | The features were computed over the MSA induced by the larger subgroup of sequences |
| 0.93% | Subgroup MSA - MSA entropy |  |
| 1.04% | Subgroup MSA - # different site-patterns |  |
| 1.83% | Subgroup MSA - % fully conserved sites |  |

**Supplementary Table S2. Complete features set.** The complete set of features that were used in the learning process. The table is divided to four groups: the model components, which were integrated as features in the training of ModelTeller and three sets of predictive features. The importance percentages signify the contribution of each feature for constructing the Random-Forest model for training ModelTeller. The features colored in red were proposed as test statistics for model adequacy or for examining models fit (the relevant references are denoted next to each). (*) Features marked with an asterisk were removed from the final version of ModelTeller due to long running times but high correlation to other features, or due to negligible effectiveness.
